# AUREOCHROME 1b is a transcriptional regulator of pigment biosynthesis in *Phaeodactylum tricornutum*

**DOI:** 10.64898/2026.06.16.732132

**Authors:** Florian Pruckner, Michele Fabris

## Abstract

Diatoms have attracted attention for their potential to produce high-value metabolites, such as terpenoids. However, the regulatory mechanisms governing their isoprenoid metabolism remain poorly understood, which poses a major bottleneck in engineering them for greater production efficiency. By mining the *Phaeodactylum tricornutum* co-regulation network (PhaeoNet), we identified two candidate transcription factors (TFs) co-regulated with the methylerythritol 4-phosphate (MEP) pathway, suggesting a potential regulatory role in isoprenoid biosynthesis: AUREOCHROME 1b (AUREO1b) and sigma factor 70.4 (σ70.4). To elucidate their mechanistic roles in isoprenoid metabolism, we generated episomal overexpression lines for both candidate TFs. These were characterized alongside lines overexpressing HSF1 and HSF3, previously identified regulators of carotenoid biosynthesis, to compare growth, pigment content, and global transcriptomic profiles. Phenotypically, overexpression of AUREO1b, PtHSF1, and PtHSF3 led to increased pigment accumulation, whereas σ70.4 overexpression did not alter pigment levels. Transcriptomic analyses revealed that each TF upregulates distinct sets of genes involved in pigment biosynthesis, pigment binding, and photosynthesis. These results expand our functional understanding of the transcriptional regulation of isoprenoid and pigment pathways in diatoms and broaden the possibilities of engineering diatom microalgae for improved production of high-value compounds.

## 1. Introduction

Marine microalgae grow in seawater using only sunlight and CO⎕, without competing for arable land, making them a production platform with great potential on sustainability [1]. Among these, the diatom *Phaeodactylum tricornutum* has emerged as a versatile model, with fast growth capacity and expanding molecular tools that enable its engineering for the synthesis of high-value compounds such as terpenoids [2–4].

Terpenoids form the largest and most structurally diverse class of natural products and derive from isoprenoids. They are built from five-carbon isopentenyl diphosphate (IPP) units, found across all domains of life [5]. Several compounds of this class play a crucial role in primary metabolism. For example, sterols maintain membrane integrity and fluidity [6], carotenoids are involved in light harvesting, protection from light stress, and defend against reactive oxygen species [7], and ubiquinones shuttle electrons [8]. In specialized metabolism, many terpenoids mediate defense and signaling [9,10]. Their potent bioactivities underlie applications in pharmaceuticals (e.g., artemisinin, taxol), nutraceuticals, flavors, fragrances, and biofuels [11].

Diatoms such as *P. tricornutum* generally possess two isoprenoid biosynthetic pathways: the plastidial methylerythritol 4-phosphate (MEP) pathway and the mevalonate (MVA) pathway, which operates mostly in the cytosol. In the chloroplast, the MEP pathway converts glyceraldehyde-3-phosphate (G3P) and pyruvate into IPP and its isomer dimethylallyl diphosphate (DMAPP) through seven enzymatic steps [12]. The enzymes 1-deoxy-D-xylulose-5-phosphate (DX) synthase (DXS) and DX reductase (DXR) catalyze the first two steps and are generally considered rate-limiting [13–15]. IPP isomerase (IDI) balances IPP/DMAPP ratios, which poses another rate-limiting step of the pathway [16]. Prenyl transferases then extend these five-carbon units into geranylgeranyl diphosphate (GGPP), which phytoene synthase (PSY) converts to phytoene, committing the metabolite to carotenoid biosynthesis [16]. In *P. tricornutum*, the major carotenoids for light-harvesting are fucoxanthin and β-carotene, while diadinoxanthin and diatoxanthin balance the redox state of the cell similar to the xanthophyll cycle [17].

The MVA pathway begins with acetyl-CoA condensation with acetoacetyl-CoA to form 3-hydroxy-3-methylglutaryl-CoA (HMG-CoA), which is further processed over four additional enzymatic steps to form IPP and DMAPP [16]. IPP is elongated to geranyl diphosphate (GPP), farnesyl diphosphate (FPP), or GGPP building blocks by prenyl transferases [18]. FPP can be committed via the alternative squalene epoxidase (AltSQE) to sterol biosynthesis [19]. Although HMG-CoA reductase (HMGR) and SQE or AltSQE often limit sterol synthesis in eukaryotes, their overexpression in diatoms does not increase sterol levels, indicating stringent regulatory control [20]. Downstream, farnesyl diphosphate (FPP) is channelled by squalene synthase (SQS) into brassicasterol, campesterol, and cholesterol biosynthesis [21].

A key challenge in boosting terpenoid yields is the cell’s homeostatic regulation. Broadly reprogramming metabolism via transcription factors (TFs) engineering and manipulation, could unlock higher production capacity [22]. In *P. tricornutum*, two heat-shock factor (HSF) TFs , HSF1 and HSF3, have been recently shown to influence carotenoid biosynthesis: the overexpression of HSF1 increases both triacylglycerol and fucoxanthin by upregulating MEP and carotenoid biosynthesis genes [23], while the overexpression of HSF3 boosts fucoxanthin accumulation through a similar activation of key carotenoid biosynthetic genes [24]. In profiling the transcriptional regulation of *P. tricornutum* isoprenoid metabolism under prolonged phosphate depletion, we identified based on the PhaeoNet co-regulation network [25] candidate transcription factors potentially involved in the regulation of the MEP pathway. These included Aureochrome 1b (AUREO1b) and Sigma factor 70.4 (σ70.4) [4]. Here we functionally characterized these candidates in diatom overexpression lines and profiled their phenotypes and transcriptomes to uncover new layers of transcriptional control over the isoprenoid metabolism. These insights expand our fundamental understanding of algal metabolism and facilitate engineering of microalgae as sustainable factories for terpenoid-based nutraceuticals, pharmaceuticals, and biofuels.

## 2. Materials and Methods

### 2.1 Cultivation, cell count measurement, and harvesting

*P. tricornutum* (strain CCMP632, CCAP 1055/1) was grown in enriched seawater artificial water (ESAW) medium [26] under continuous light, with a light intensity of 90 µmol photons m⎕² s⎕¹. Cultures were kept at 21 °C in an Innova S44i orbital shaking incubator (Eppendorf, Germany) with constant agitation at 95 rpm. Cell density was measured on a Guava H5 flow cytometer (Cytek, Japan), diluting samples 1:10 prior to analysis to prevent detector saturation.

On the last day of the experiment, 10^9^ cells were harvested by centrifuging the calculated volume of cell culture for 2 min at 4200 RCF, resuspending the pellet in 1 ml of PBS, transferring the resuspension into 2 ml microcentrifuge tubes, centrifuging for 1 min at 17000 RCF, removing the supernatant, and flash freezing the pellet in liquid nitrogen, and storing until extraction at -70 °C.

### 2.2 Episomes assembly and diatom conjugation

Episomal vectors were constructed with the universal loop (*u*Loop) assembly system [27]. In brief, each coding sequence was retrieved from Ensembl [28] (*AUREO1b*: *Phatr3_J15977.t1*; σ*70.4*: *Phatr3_J9312.t1*; *HSF1*: *Phatr3_J49566.t1*; *HSF3: Phatr3_J44200.t1*). The sequences were domesticated with removal of Sap-I and Bsa-I restriction sites using Benchling [29], synthesized by Azenta, GENEWIZ (Germany), and cloned into “CD” L0 parts. The L0 parts were combined with the *Phatr3_J49202* promoter region (“AC”), a 3×Ostop cassette (“DE”), and the *Phatr3_J25172* (FcpB) terminator region (“EF”) into a pCAo-4 receiver backbone. A pCAo-2 fluorescent reporter cassette (mVenus, flanked by the same “AC”, “DE”, and “EF” elements) was similarly created and used in the assembly for tracking the episome presence and function in diatoms via flow cytometry. Along with a previously built pCAo-1 plasmid, carrying the “*Sh-ble*” Zeocin resistance gene, the yeast “*CEN-ARS-HIS*” maintenance region, and a pCAo-3 carrying a spacer sequence, pCAo-1, 2, 3, and 4 were assembled into a pCAe-1 receiver vector, yielding the final episomes. Sequences of the L2 episomes are provided in Supplementary Material (File S1-S5).

Episomes were introduced into *P. tricornutum* via bacterial conjugation with a protocol adapted from Karas et al. (2015) [30]. An exponentially growing *P. tricornutum* culture was used to inoculate a 50 mL culture with a starting optical density at 750 nm (OD_750_) of ∼0.03. The culture was grown until it reached OD_750_ of 0.3. Six well plates were prepared to contain ∼3.5 mL of ½ ESAW–5% Luria Bertani (LB)–1% agar per well and left under the laminar flow hood for 1 h to dry completely. The day before conjugation, *P. tricornutum* cultures in the mid to late exponential growth stage (OD_750_ =O0.3) were centrifuged at 3000 × g. For every 50 mL of harvested culture, 500 μL of ESAW was added, and 50 μL of this mixture was spotted onto the sixOwell ½ESAW–5%LB agar plates. The plates were allowed to dry in a laminar flow hood until no visible liquid remained. Plates were then incubated under continuous light for ∼18 h at 90Oμmol photons · m−2· s−1 at 21°C.

An overnight culture of an *Escherichia coli* Epi300 strain containing the conjugative plasmid (pTAOMOB) and the cargo plasmid (episome) was started the night before the conjugation experiment (shaking at 180 rpm, 37°C). The overnight culture was used to inoculate 20 mL of fresh *E. coli* culture with a 1:50 dilution in LB supplemented with gentamicin 20 μg · mL−1 and spectinomycin 50 μg · mL−1 . The flasks were shaken at 200 rpm and 37°C until the culture reached an OD600 of 0.8–1.0. The culture was then spun down for 10 min at 3000 × g. All supernatant was removed, and the cell pellet was gently resuspended in 250 μL of super optimal broth with catabolite repression (SOC) medium. Fifty microliters of the *E. coli* culture were pipetted onto the dried diatom spot in the multi well plate. The spot was not spread but allowed to dry under laminar flow. The plates were incubated for 90 min at 30°C in the dark and then moved to diatom growth conditions for a 3 day recovery period. After 3 days, 1 mL of ESAW medium was added to the ½ESAW/5%LB plates. Cells were resuspended and transferred to a ½ESAW agar plate with 100 μg · mL−1 zeocin and a suitable antibiotic against bacteria to select for transformed diatoms. Colonies of *P. tricornutum* were picked after 2–3 weeks using a sterile pipette tip and resuspended in 200 μL ESAW and 100 μg · mL−1 zeocin medium in 96 well plates. Diatom exconjugants were screened on a Guava easyCyte HT flow cytometer (488 nm excitation, FITC detection). Four independent cell lines per TF were selected based on mVenus fluorescence intensity and population uniformity and subsequently used for physiological and molecular characterization.

### 2.3 Pigment extraction and quantification

Frozen samples with wet cell pellets were kept on metal blocks precooled with liquid nitrogen, and were freeze-dried in a Martin Christ ALPHA 1–2 LDplus vacuum chamber overnight. Pigments were extracted from each dried pellet by adding 1.5 mL of methanol (95% v/v) buffered with 2% ammonium acetate (w/v), manually dissolving of the pellet with pointy tweezers, and incubating in the dark at 4 °C on a rotating mixer for three hours. The extract was clarified by centrifugation at 12000 × g for five minutes at –4 °C, and 1 ml of the supernatant was transferred into amber HPLC vials. Pigments were separated on an Agilent 1200 Series HPLC system equipped with an autosampler, column thermostat, and diode array detector, using a YMC Carotenoid column (CT99S03-1503WT). A binary solvent system of acetone/methanol (40:60, solvent A) and water/acetone (40:60, solvent B) was delivered at 0.5 mL minO¹ with a 5 µL injection volume and a column temperature of 40 °C. The gradient commenced at 40% solvent A, rose to 70% over three minutes and held until 22 minutes, increased to 90% by 26 minutes with a five-and-a-half-minute hold, then returned to 60% over 3.5 minutes for a total run time of 55 minutes. Pigment peaks were monitored at 450 nm.

### 2.4 RNA extraction, RNA sequencing, and data processing

RNA extraction and sequencing was performed as previously described in [4].

### 2.5 Quantitative real-time PCR transcript quantification

RNA extraction from *P. tricornutum* and cDNA synthesis was done as described previously [4]. Quantitative real-time PCR (RT-qPCR) was performed using FastStart Essential DNA Green Master (Roche, #06924204001) on a LightCycler 480 Instrument II (Roche). Reactions were run with an initial pre-incubation/polymerase activation step at 95 °C for 5–10 min, followed by 45 cycles of denaturation at 95 °C, primer annealing at 55 °C, and extension at 72 °C, with a single fluorescence acquisition at the end of each elongation step. Relative gene expression was calculated using the comparative Ct method and normalized to the reference gene β-tubulin (TubB; *Phatr3_J21122*). All qPCR primers used in this study are listed in Table S1.

### 2.6 Gene ontology analysis

For gene ontology molecular function enrichment, we processed lists of differentially expressed genes (DEGs) through an automated R pipeline (see File S6). Briefly, each gene list was stored as a tab delimited text file, and was read into R. GO annotations specific to *P. tricornutum* were then retrieved from the Ensembl Protists database, and GO identifier and their descriptive terms were linked to each gene. Enrichment testing was performed using the “clusterProfiler” function enricher(), with *p* values adjusted by the Benjamini–Hochberg method, and a *q* value threshold of 0.05. The top 15 GO terms ranked by fold enrichment were selected for visualization.

### 2.7 Phylogenetic analysis

Homologues of the AUREO1b, HSF1, and HSF3 were retrieved, by sequence homology search of the protein sequence as defined on Ensembl protists [28] using BLASTp [31] (E-value < 0.01). To avoid hits based only on small, conserved domains, we considered only hits with more than 75% query coverage. The hits were exported in fasta file format, and uploaded and analyzed on NGPhylogeny.fr [32]. Tree graphics were formatted using iTOL [33].

## 3. Results and Discussion

### 3.1 Genes encoding candidate transcription factors co-regulate with isoprenoid biosynthesis pathways

Using the PhaeoNet dataset [25], we previously showed that within the terpenoid metabolism especially the MEP pathway has a high degree of co-regulation within the lightsteelblue1 module [4]. Strikingly, the lightsteelblue1 module included key nodes of the MEP pathway, such as DXS, DXR, and IDI, in addition to 4-hydroxy-3-methylbut-2-enyl diphosphate synthase (HDS) and 4-hydroxy-3-methylbut-2-enyl diphosphate reductase (HDR), representing the last two steps of the pathway. As also shown in our previous work [4], three TFs are co-regulated with these genes, within the lightsteelblue1 module: *Aureo1b* (*Phatr3_J15977*), σ*70.4* (*Sigma70.4*, *Phatr3_J9312*), and *Phatr3_J50411*, which hence are prime TF candidates to regulate the MEP pathway (Fig. 1).

**Figure 1.**
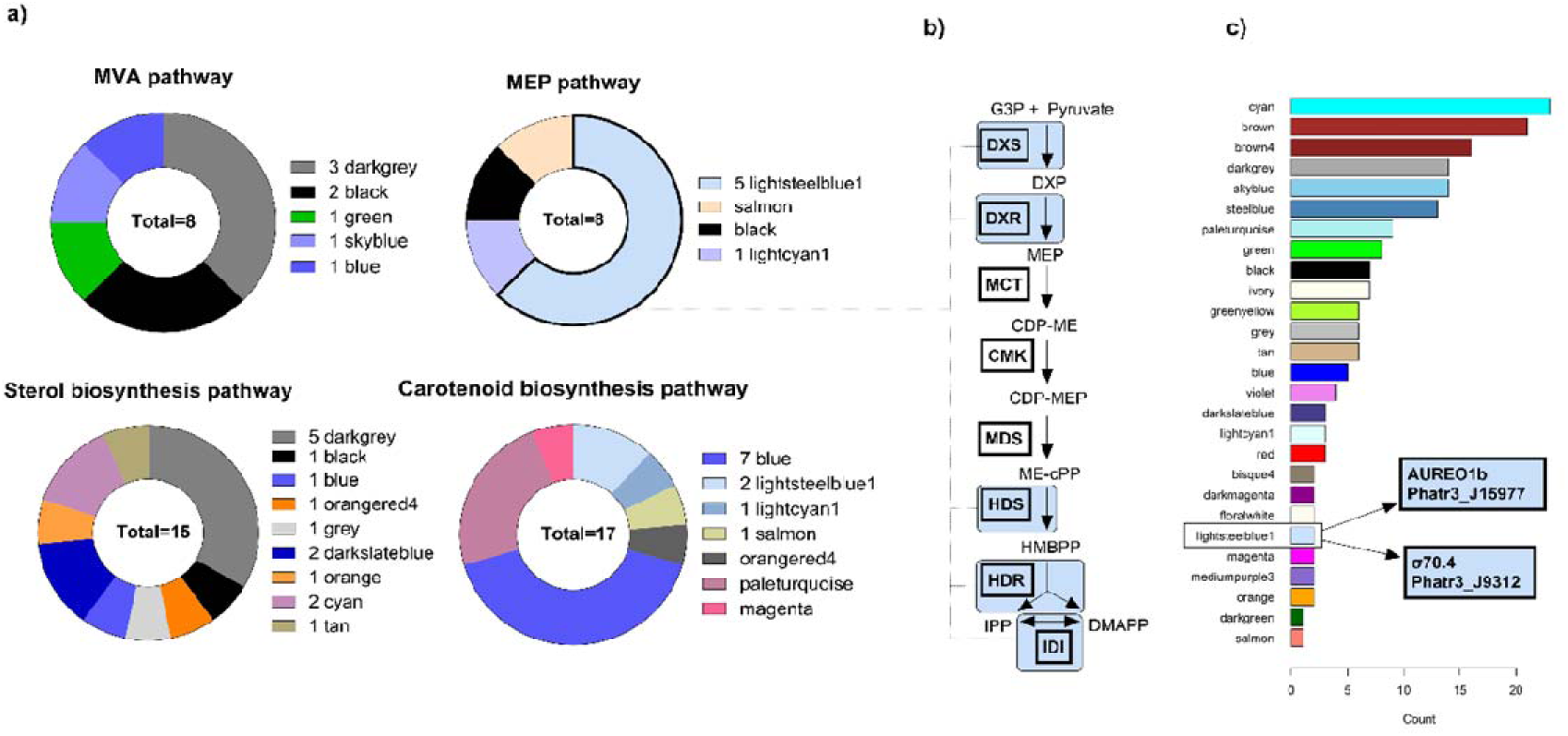
Co-regulation analysis of genes involved in the isoprenoid metabolism in PhaeoNet. **(a)** Distribution of terpenoid-biosynthesis genes across PhaeoNet co-expression modules, with each module color indicating a distinct cluster. **(b)** Detailed view of the MEP pathway genes, which cluster predominantly within the lightsteelblue1 module (shaded in blue). **(c)** Mapping of transcription factors of Rayko *et al*., 2010 [53] onto PhaeoNet modules, with the two lightsteelblue1 TFs, AUREO1b and σ70.4, highlighted as putative regulators of isoprenoid metabolism. **Abbreviations:** DXS, 1-deoxy-D-xylulose 5-phosphate synthase; DXR, 1-deoxy-D-xylulose 5-phosphate reductoisomerase; MCT, 2-C-methyl-D-erythritol 4-phosphate cytidylyltransferase; CMK, 4-cytidine-5-diphospho-2-C-methyl-D-erythritol kinase; MDS, 2-C-methyl-D-erythritol 2,4-cyclodiphosphate synthase; HDS, 4-hydroxy-3-methylbut-2-enyl diphosphate synthase; HDR, 4-hydroxy-3-methylbut-2-enyl diphosphate reductase; IDI, isopentenyl diphosphate isomerase.

We overexpressed the TFs AUREO1b and σ70.4 in *P. tricornutum* to assess their impact on pigment accumulation and to profile global transcriptomic changes. An overexpression line for the third candidate TF, Phatr3_J50411, was also created, but was ultimately not included in the analysis of this study, as this line was lost due to contaminations. Additionally, we generated overexpression lines also for HSF1 and HSF3, previously shown to enhance pigment content [23,24], to serve both as positive controls and to extend RNA-seq analysis to these TFs, which have not yet been characterized at the transcriptome level.

Aureochrome TFs (AUREOs) are bZIP TFs that can sense and react to light via their light-oxygen-voltage (LOV) domain [34]. They are found in diatoms and can facilitate rapid transcriptional responses, by altering their DNA-binding capacity directly, depending on conformational changes induced by light [34]. *P. tricornutum* encodes four AUREOs; AUREO1a orchestrates blue-light–dependent transcription and non-photochemical quenching [35–39], while AUREO1c drives rapid non-photochemical quenching (NPQ) induction under high-light stress [40]. Although AUREO1b has been linked to blue-light adaptation, its role in carotenoid regulation remains unexplored [41].

The σ70 family, originally derived from bacteria, is associated with the regulation of plastid-encoded genes of the σ70 family [42]. Nucleus-encoded sigma factors are thought to be imported into the plastid, where they guide the plastid RNA polymerase in the transcriptional regulation of chloroplast genes [25,43]. The *P. tricornutum* genome encodes eight distinct σ-factors, of which only three possess predicted chloroplast-targeting peptides, while the remaining lack apparent transit signals and may perform alternative or indirect regulatory functions [25]. σ70.4 (Phatr3_J9312), which is co-regulated with the MEP pathway in *P. tricornutum*, belongs to this latter group, as it is not predicted to localize to the chloroplast [25]. However σ70.4 may still influence plastid-associated transcript levels indirectly over other TFs, or through modulation of nucleus-encoded plastid genes, such as those of the MEP pathway.

### 3.2 The overexpression of AUREO1b, HSF1, and HSF3 enhance pigments accumulation, while σ70.4 does not

To evaluate the role of the identified candidate TFs on pigment biosynthesis and growth, we phenotyped *P. tricornutum* overexpression (OE) lines for AUREO1b, σ70.4, HSF1, and HSF3, compared to diatom cell lines expressing mVenus only, as negative control (Fig. 2). Growth curves revealed that σ70.4QOE lines grew comparably to the controls, whereas HSF1Q and HSF3QOE lines displayed a modest reduction in growth (Fig. 2a). AUREO1bQOE lines showed slowest growth, with the greatest difference in cell density to control lines by day 8 (Fig. 2b).

**Figure 2.**
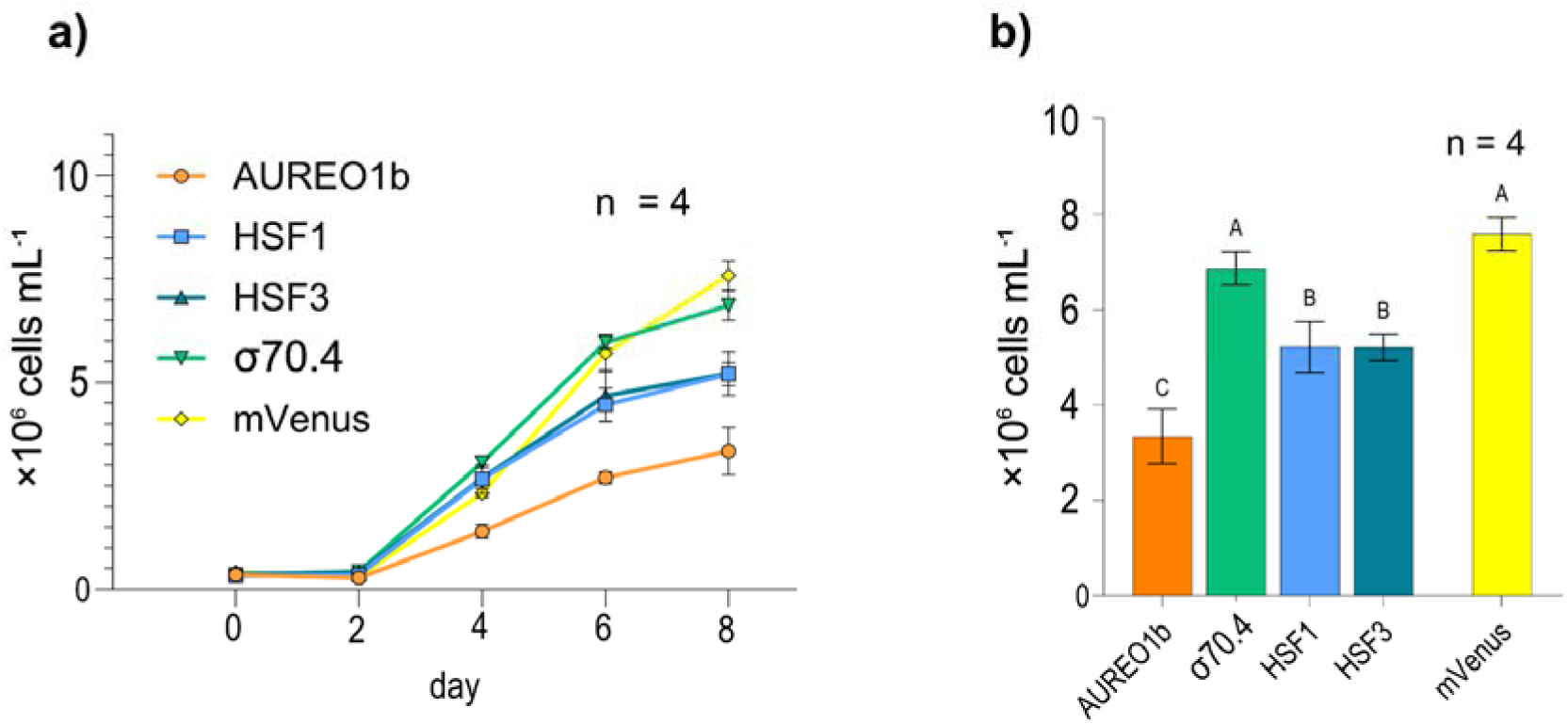
Growth of transgenic diatom lines overexpressing TFs. **(a)** Growth curves of four TF-OE lines (AUREO1b-OE, σ70.4-OE, PtHSF1-OE, PtHSF3-OE) and mVenus as a control line, monitored over eight days. **(b)** Cell density on day 8 for each TF-OE line. Different letters above bars indicate statistically significant differences (one-way ANOVA, p-value < 0.01, n=4). Error bars indicate the standard deviation.

Notably, earlier studies reported that HSF1- and HSF3-OE lines grew identically to WT under a 16:8 h light:dark regime at 50 μmol photons mO² sO¹ [23,24], whereas our HSF1-OE and HSF3-OE lines, grown under continuous light at 90 μmol photons mO² sO¹, showed diminished growth. This may relate to the light-dependent expression of these TFs. HSF transcript levels in *P. tricornutum* are strongly light-regulated, with HSF3 in particular showing clear light-dependent expression [44]. Consistent with this, transcriptomic data comparing *P. tricornutum* under continuous light versus a 12:12 h light:dark regime showed that HSF1 and HSF3 are differentially expressed during the dark phase, but in opposite directions [45]: at 1h (ZT13) and 7h (ZT19) after onset of dark, HSF1 was upregulated (fold-change 1.65 and 1.61; *p* = 0.015 and 0.029) and HSF3 downregulated (fold-change 0.60 and 0.48; *p* < 0.01 for both) (Supplementary Data S4; unpaired two-tailed *t*-test). Thus, although overexpression of either factor alone yields a similar high-pigment phenotype, both appear to play light-dependent roles in the broader regulation of cell physiology. This could indicate that slower growth of HSF-OE lines observed in this study but not in previous ones, may be attributable to the continuous light conditions.

After 8 days of growth, biomass was harvested for pigment analysis, which revealed comparable increases in carotenoid and chlorophyll accumulation in AUREO1bO, HSF1-, and HSF3OOE lines relative to controls, while σ70.4OOE lines showed no detectable changes (ANOVA, *p*-value < 0.01) (Fig. 3). This confirms the increased accumulation of fucoxanthin previously reported in HSF1- [23] and HSF3-OE lines [24]. The changes in carotenoids fit with previous studies which identified AUREO1b as a potential low-light associated transcription factor [37,41]. However, *PtAUREO1b* knockout mutants grown under a 14/10 h blue light regime also show elevated chlorophyll *a* levels—similar to the OE lines in this study [41]. Notably, this phenotype was not observed under red light in the same study [41]. This suggests that chlorophyll *a* levels are regulated by AUREO1b in a light quality–dependent manner, while AUREO1b expression is also regulated by light/dark cycles [46], complicating interpretation of why overexpression under continuous white light in this study produces a phenotype similar to that of knockout under blue light. Hence it would be interesting to investigate the phenotype of AUREO1b knockout lines in direct comparison with overexpression lines under different light conditions, with a special focus on carotenoid levels, which have not been subject of investigation in previous studies around AUREO1b [37,41,46]. To elucidate the transcriptional networks underlying these phenotypes, we extracted total RNA from day 8 biomass and performed RNA-seq analysis.

**Figure 3.**
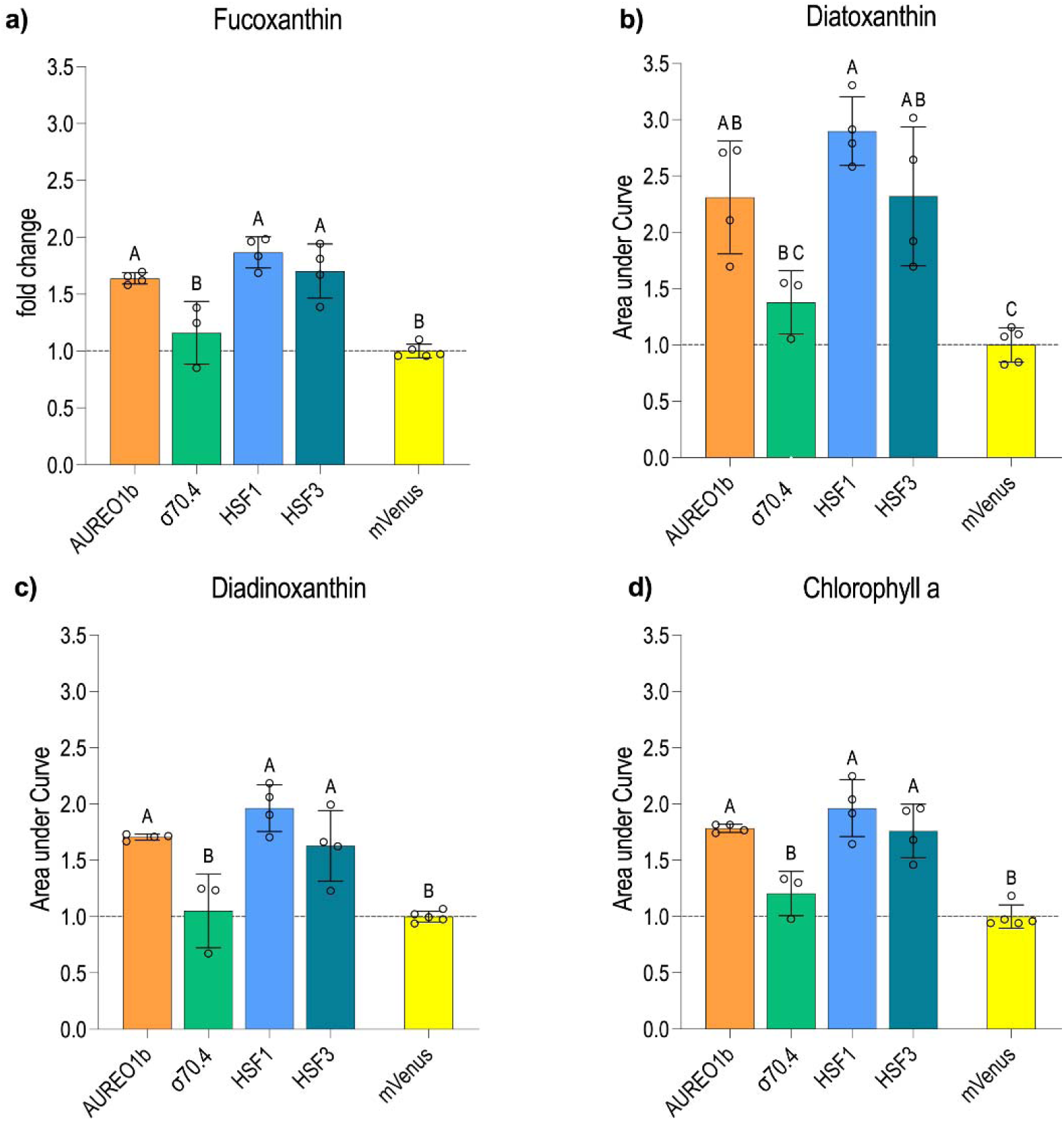
Relative changes in pigment accumulation in TF-OE cell lines. Quantification of **(a)** fucoxanthin, **(b)** diatoxanthin, **(c)** diadinoxanthin, and **(d)** chlorophyll a. For each sample pigments were extracted from 10a cells, and the area under curve was directly measured from the chromatograms, without any additional normalization. Each TF-OE line was analyzed with four biological replicates, while five replicates were used for the mVenus OE control cell lines. Significant differences between groups were determined by one-way ANOVA (p < 0.01) and are indicated by different letters above the bars. For groups showing significant differences relative to the mVenus control, the change is expressed as a percentage relative to the control line. Error bars indicate the standard deviation.

### 3.3 Overexpression of AUREO1b, σ70.4, HSF1, and HSF3 induces pigment- and photosynthesis-related gene expression

To investigate whether the observed phenotypic changes in pigment levels are an effect of carotenoid and chlorophyll biosynthesis pathways, and whether lightsteelblue1 TFs indeed promote MEP pathway expression levels, we performed RNA-seq and gene expression analyses on RNA isolated from biomass samples of the aforementioned experiment for each TF-OE line (initially 4 biological replicates per OE, 5 for mVenus control). After removing low-quality libraries, the final replicate counts were n=3 for AUREO1b, n=4 for HSF1, n=2 for HSF3, and n=3 for mVenus. Principal component analysis (PCA) confirmed distinct clustering of replicates in groups (Fig. S1a). Differential expression analysis (|log FC| > 1, adjusted *p*-value < 0.01) revealed 1411 differentially expressed genes (DEGs) in AUREO1b-OE, 803 in σ70.4-OE, 1694 in HSF1-OE, and 1059 in HSF3-OE lines (Tables S2–S5). Overlap of DEGs is shown in a Venn diagram in Fig. S1b. As a validation of the experiment we evaluated transcript levels of overexpressed TF transcript in the cDNA used for sequencing via quantitative real time PCR (qRT-PCR), which confirmed that TF transcripts were upregulated in the respective overexpression line (Fig. S2). When evaluating the change in expression levels of the 4 TFs in our transcriptomic datasets we observed similar upregulation of the TF overexpressed in the respective line, with no upregulation of non-overexpressed TFs, indicating that these TFs do not influence each other’s transcript levels (Fig. S2b).

Gene ontology (GO) analysis of differentially expressed genes (DEGs) returned a set of modestly enriched categories (<4.5-fold change) but did not point to a single dominant molecular function (Fig. S3). Because GO definitions are often broad or unspecific, we manually inspected key metabolic pathways (gene lists in Table S6) and found a clear, consistent signal: all four TF-OE lines increased expression of genes linked to pigment biosynthesis and pigment-binding complexes, including components of the MEP pathway, carotenoid and chlorophyll biosynthesis, light-harvesting complexes (LHCs), photosystem 2 (PSII) subunits and other electron-transport components (Fig. 4).

**Figure 4.**
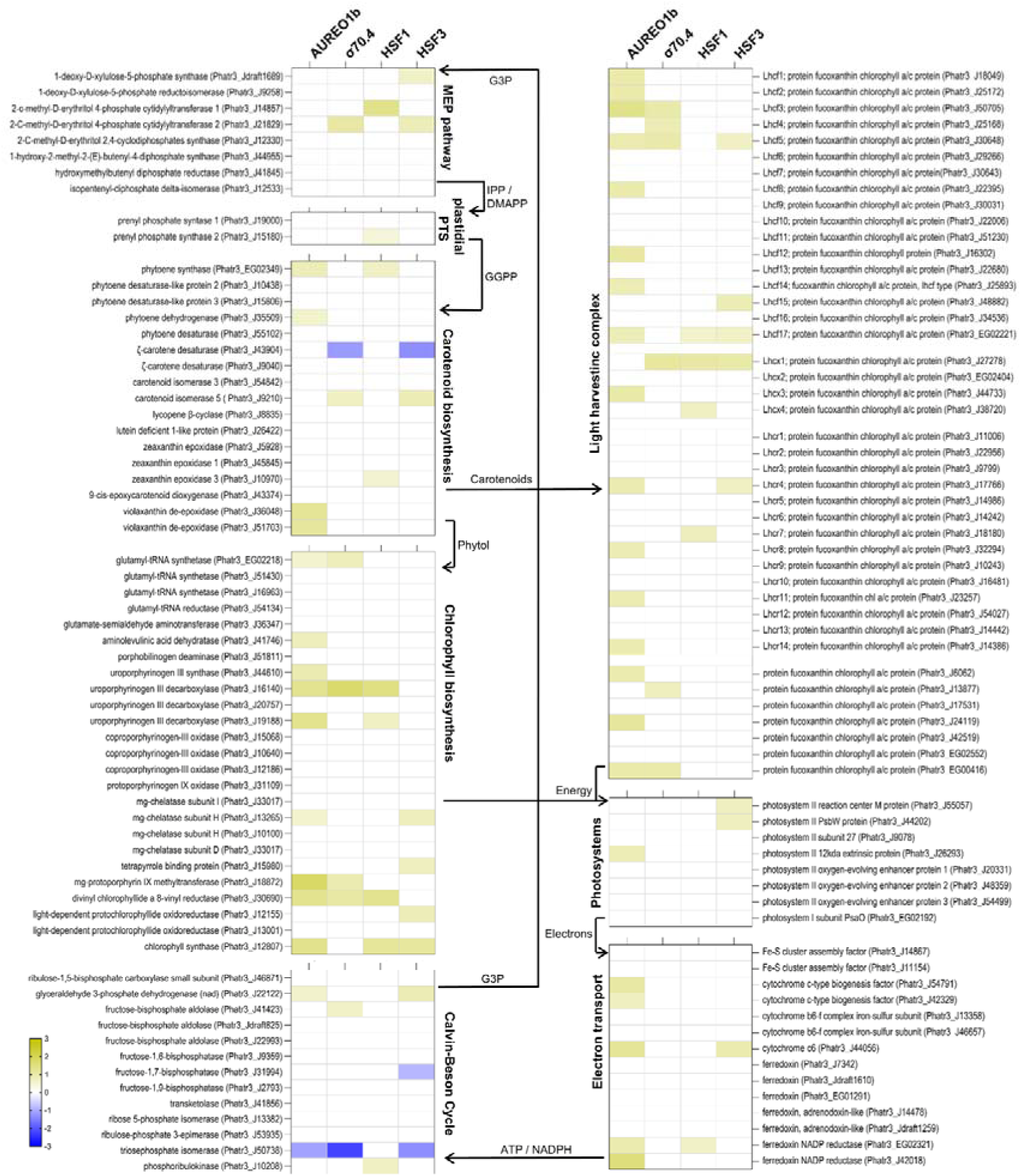
Heatmap of differential gene expression in pigment biosynthesis, pigment-binding, and photosynthesis pathways across transcription factor overexpressing (TF-OE) lines. Rows represent individual genes, grouped by pathway, with arrows indicating the dependence between these processes. Columns correspond to TF-OE lines AUREO1b-OE, σ70.4-OE, PtHSF1-OE, and PtHSF3-OE. Cell color indicates loga fold-change relative to the mVenus control (blue = downregulation; yellow = upregulation).

Despite selecting AUREO1b for its co-expression with MEP genes, AUREO1b-OE lines did not differentially regulate genes involved in the initial MEP pathway cluster in our experiments. Similarly, in σ70.4-OE cell lines, only the *2-C-methyl-D-erythritol 4-phosphate cytidylyltransferase 2* (*MCT2*; *Phatr3_J21829)* was upregulated, a gene not previously observed as co-regulated with σ70.4 in PhaeoNet [25]. This suggests that the MEP lightsteelblue1 cluster, AUREO1b and σ70.4 might be controlled by a common upstream regulator rather than either of the TFs controlling the MEP pathway. Overexpression of *HSF3* specifically induced *DXS*, as has been reported in earlier work [24]. In both σ70.4- and HSF3-OE cell lines, a *carotenoid isomerase 5* (*CRTISO5*; *Phatr3_J9210*) was upregulated, while one ζ*-carotene desaturase* (*Phatr3_J43904*) was downregulated. AUREO1b- and HSF1-OE both induced *Phatr3_EG02349* (*PSY*), which is in line with HSF1 having been shown to upregulate *PSY* [23]. AUREO1b additionally upregulated two violaxanthin de-epoxidases (*Phatr3_J36948* and *Phatr3_J51703*), which suggest the occurrence of a possible rebalancing mechanism from diadinoxanthin to diatoxanthin. This aligns with measured pigment changes (diatoxanthin +131% vs diadinoxanthin +70%) (Fig. 4). However, HSF1-OE lines also showed a shift towards an increased diatoxanthin:diadinoxanthin ratio even though zeaxanthin epoxidase 3 (*ZEP3*; *Phatr3_J10970*) was upregulated rather than VDEs.

Carotenoids are incorporated into LHCs where they serve light-harvesting (mainly LHCf and LHCr classes) or photoprotective roles (LHCx class). All TF-OE cell lines upregulated subsets of the 42 annotated LHC genes, with differing patterns (Fig. 4). Sixteen genes were upregulated in AUREO1b-, six in σ70.4-, four in HSF1- and five in HSF3-OE lines. Although upregulated genes spanned several LHC classes, distinct patterns emerged: σ70.4, HSF1 and HSF3 induced *Phatr3_J27278* (LHCx1), a gene critical for high-light acclimation [47], whereas AUREO1b did not. HSF1 induced LHCx4 while AUREO1b specifically induced LHCx3, which has been associated with NPQ induction and blue-light responses [48], consistent with AUREO1b’s role as a blue-light photoreceptor. AUREO1b also upregulated four LHCr genes (photosystem 1 PSI-associated), while HSF1 and HSF3 each induced one and σ70.4 induced none. This upregulation of LHCr transcripts coincided with upregulation of the cyclic electron flow (CEF) regulator *Phatr3_J44748* (PGR5) in AUREO1b (log2FC = 1.38), HSF1 (log2FC = 1.04) and HSF3 (log2FC = 0.93), but not in σ70.4.

Chlorophyll biosynthesis genes were broadly induced across all OE lines, although with TF-specific patterns. AUREO1b, σ70.4 and HSF1 upregulated *uroporphyrinogen III decarboxylase (Phatr3_J16140*) and *divinyl chlorophyllide a 8-vinyl reductase* (*Phatr3_J30690*). *Chlorophyll synthase* (*Phatr3_J12807*) was induced by AUREO1b, HSF1 and HSF3. *Glutamyl-tRNA synthase* (*Phatr3_EG02218*) and *Mg-protoporphyrin IX methyltransferase* (*Phatr3_J18872*) were induced by AUREO1b and σ70.4. Uroporphyrinogen decarboxylase (*Phatr3_J19188*) was specifically upregulated in AUREO1b-and HSF1-OE lines. Although overexpression of single chlorophyll biosynthesis enzymes, such as for example *chlorophyll synthase*, can sometimes increase chlorophyll levels in plants [49], the physiological effect of upregulation of individual enzymes in diatoms remains unclear. Nevertheless, the transcriptional pattern – upregulation in AUREO1b, HSF1 and HSF3 but not in σ70.4 – matches the observed higher chlorophyll accumulation in those three lines (Fig. 4).

Changes in the expression of genes encoding photosystem subunits were modest and TF-dependent. σ70.4- and HSF1-OE lines showed no differential regulation of any PS subunit; HSF3 induced *PSII reaction center M* (*Phatr3_J55057*) and *photosystem II reaction center protein W (PsbW; Phatr3_J44202)*, while AUREO1b upregulated *PSII 12 kDa extrinsic protein (Phatr3_J26293*). Although homologues of these proteins are important for photosystem function in plants and cyanobacteria [50,51], functional data in diatoms are limited, so their physiological roles here require further validation.

Several electron transport components were particularly induced in AUREO1b lines, including cytochrome c-type biogenesis factors (*Phatr3_J54791*, *Phatr3_J42329*), cytochrome c6 (*Phatr3_J44056)* and ferredoxin-NADP reductases (*Phatr3_EG02321*, *Phatr3_J42018*). HSF1-OE induced *Phatr3_EG02321* while HSF3-OE induced *Phatr3_J44056*. This pattern could suggest a possibly stronger capacity for electron transfer and NADPH generation in AUREO1b OE-lines. This hypothesis is supported by the fact that photosynthesis rates were reported to drop in AUREO1b knockout (KO) lines [41].

In the Calvin–Benson cycle, AUREO1b- and HSF3-OE upregulated *G3P dehydrogenase* (*Phatr3_J22122*), σ70.4 induced *fructose-bisphosphate aldolase* (*Phatr3_J41423*), and HSF3 induced *phosphoribulokinase* (*Phatr3_J10208*). Notably, AUREO1b-, σ70.4- and HSF3-OE all led to downregulation of *triosephosphate isomerase* (*Phatr3_J50738*), which could shift the balance between dihydroxyacetone phosphate (DHAP) and G3P. In AUREO1b- and HSF3-OE lines, where *G3P dehydrogenase* is also upregulated, this may lead to G3P accumulation and redirection of carbon toward prenyl phosphate or lipid biosynthesis.

Although AUREO1b-, σ70.4- and other HSF-OE lines share overlapping expression signatures, the phenotype of σ70.4-OE is not associated with increased pigment accumulation (Fig. 3) despite some similar transcriptional changes (Fig. 4). The extensive interconnection among isoprenoid and photosynthetic pathways, with MEP-derived IPP/DMAPP feeding carotenoid synthesis, carotenoids and phytol integrating into LHCs, which in turn drive electron transport, and generate ATP and NADPH for the Calvin–Benson cycle, which ultimately regenerates MEP precursors (Fig. 4), complicates the explanation of the lack of an increased pigment phenotype in σ70.4_OE cell lines.

Comparison of our HSF1 and HSF3 transcriptional profiles with previous reports shows only partial overlap. HSF1-OE upregulated PSY and induced ZEP3 in our data (whereas earlier work reported ZEP1), and we did not observe the previously reported upregulation of fatty-acid biosynthesis in HSF1-OE [23]. In this study HSF3-OE induced key carotenoid biosynthesis genes, such as *DXS* and *CRTISO5*, but not the broader carotenoid biosynthesis gene set reported previously [24]. Differences in light regime between studies (continuous 90 µmol m⎕² s⎕¹ here versus 16:8 h at 50 µmol m⎕² s⎕¹ previously [23,24]) likely account for at least part of these discrepancies. Overall, the most robust trend across all TF-OE lines was coordinated upregulation of genes linked to pigment biosynthesis and pigment binding, covering carotenoid and chlorophyll pathways as well as LHC genes, which aligns with the altered pigment phenotype observed in AUREO1b, HSF1 and HSF3 OE cell lines (Fig. 4).

The contrast between σ70.4 and AUREO1b is nonetheless informative: despite sharing overlapping transcriptional patterns, only AUREO1b overexpression translates into detectable pigment accumulation, reinforcing that transcriptional upregulation of biosynthetic genes is necessary but likely not sufficient for phenotypic change. This could reflect the presence of other layers of regulation, for example post-translational, which was not captured by our analyses and remains to be determined.

### 3.4 AUREO1b and HSF3 are conserved among diatoms, while HSF1 is *P. tricornutum* specific

To assess the phylogenetic distribution of TFs that were shown to boost carotenoid biosynthesis in this study, we submitted protein sequences of AUREO1b, HSF1, and HSF3, to BLASTp searches [31] using an E-value threshold of 1 ×1 0⁻² and required ≥75% query coverage to minimize matches driven only by conserved domains (e.g., the HSF DNA-binding domain or the LOV domain). Under these criteria, we identified 1 9 homologues of AUREO1 b and 45 homologues of HSF3, all in diatom species, with no hits recovered from non-diatom lineages (Fig. S4). For HSF1, no homologues meeting our coverage threshold were detected in any species. Although HSF1-like proteins have been reported in plants and green algae in another study [23], no query coverage thresholds have been applied in that study, yielding homologues likely based on sole homology of the conserved HSF DNA-binding domain. Accordingly, in this analysis HSF1 appears *P. tricornutum*-specific, whereas AUREO1b and HSF3 are conserved in diatoms. It is therefore possible that AUREO1b and HSF3 homologues could exhibit similar roles in other diatom species.

## 4. Conclusions

In this study, we set out to identify candidate TFs that coordinate the biosynthesis of isoprenoids in the model diatom *P. tricornutum*. By integrating co-expression network analysis (PhaeoNet modules [25]) with previously composed lists of TFs [52], we identified two novel candidates that might influence carotenoid biosynthesis: AUREO1b, which was previously shown to be light-responsive and was linked to photoacclimation [41,46], and σ70.4. We overexpressed these TFs, along with the previously described carotenoid accumulation influencing TFs HSF1 and HSF3, in *P. tricornutum*, and evaluated the pigment accumulation of those lines, and conducted RNA-seq experiment to evaluate their regulatory function in detail. AUREO1b, HSF1, and HSF3 overexpression drove significant increases in both carotenoid and chlorophyll accumulation, and showed upregulation of related genes. σ70.4 overexpression, despite showing a similar upregulation of pigment- and photosynthesis-related genes on a transcriptome level, failed to elevate pigment content. RNA-seq profiling confirmed that all four TFs enhance expression of key carotenoid and chlorophyll biosynthesis, light-harvesting complex genes, illustrating a shared regulatory signature that nonetheless can produce distinct phenotypes.

Even though our approach of using co-regulation of TFs with key enzymes of the MEP pathway to find TFs that increase carotenoid content indeed led us to discover the TF with the desired phenotype (AUREO1b), MEP pathway gene expression remained markedly unaffected by the expression of this TF. This may indicated a yet undiscovered TF regulating the expression of both the MEP pathway genes and AUREO1b. σ70.4 overexpression similarly did not influence any of the genes reported to be co-regulated with it. The identification of these elusive regulators of the MEP pathway and via additional regulation of TFs such as for instance AUREO1b, the carotenoid biosynthesis pathway could allow to increase the metabolic flux through this pathway. Importantly however, in plants the MEP pathway is not only controlled on a transcriptional level by TFs [53], but also post-translationally via enzyme degradation [54], metabolite allosteric feedback inhibition [55], or indirectly via cross-talk with the cytosolic mevalonate (MVA) pathway [56,57,58,59]. This complexity of post-translational MEP pathway regulation in plants underlies the lack of understanding how diatoms control this pathway post-transcriptionally. Such post-transcriptional regulation layers may explain how MEP pathway flux may differ while transcript levels remain unaltered.

Our work was conducted under continuous high-light conditions (90 µmol photons m⎕² s⎕¹), which differ from the 16:8 h light:dark regimes at 50 µmol photons m⎕² s⎕¹, used in previous studies on HSF1 [23] and HSF3 [24]. Light regimes, quality, and intensity are well known to influence pigment biosynthesis in microalgae [58]. This discrepancy might be the explanation for the partial overlap of pigment biosynthesis genes regulation pattern observed here and in the initial studies to be differentially regulated by HSF1 and HSF3. Also, it must be kept in mind that variable replicate numbers, particularly the two replicates for HSF3, reduce statistical power, making it likely that some DEGs could not be detected. Finally, without direct TF promoter binding assays (e.g., ChIP-seq), we could not conclusively distinguish primary targets from indirect downstream effects. Implementing such studies will be crucial in further understanding how the TFs described in this study exerted their regulatory function.

The potential tight interdependency of light regimes and the detectable phenotypic effect of TF overexpression is a gap we did not explore in this study. Given the strong influence of light on carotenoid and chlorophyll biosynthesis and photosynthesis, it is likely that some phenotypes become only apparent in special light conditions, such as high- or low-light, fluctuating light, photoperiod, or light quality. Experimenting with light conditions could for instance also unlock a phenotypic response from σ70.4-OE lines.

Finally, simultaneous co-overexpression of multiple TFs may yield synergistic increases in flux through the MEP pathway and carotenoid biosynthesis. Coupling such TF combinations with the introduction of a heterologous isoprenoid synthesis pathway in the chloroplast could further amplify terpenoid production, positioning diatoms as robust, sustainable platforms for industrial biosynthesis.

## 5. Author Contribution

MF and FP conceptualized the study. FP performed experiments. FP conducted the RNA-seq data analysis. FP wrote the original draft of the manuscript and made the figures. MF reviewed and edited the manuscript. MF acquired the funding, supervised, and administered the project. All authors read and approved the final version of the manuscript.

## 6. Acknowledgements

RNA sequencing was carried out at the Center for Functional Genomics and Tissue Plasticity (ATLAS) in the Functional Genomics & Metabolism Research Unit of the University of Southern Denmark. The authors thank Gimel Pucci Infante Jørgensen and Ronni Nielsen (ORCID: 0000-0001-6331-7660) for sequencing assistance. This work, FP and MF were supported by Villum Fonden (grant number 37521 to MF) and the authors acknowledge support from the SDU Climate Cluster (SCC) with a Research Infrastructure Grant to MF.

## 7. Supplementary Materials

**Supplementary Fig. S1:** Quality control and differential expression overview of RNA-seq data.png

**Supplementary Fig. S2:** Validation of transcription factor overexpression.png

**Supplementary Fig. S3:** GO analysis of DEGs of transcription factor overexpressing lines.png

**Supplementary Fig. S4:** Phylogenetic analysis of TFs that influence pigment accumulation.png

**Supplementary Table S1:** Primers used for qPCR quantification.xlsx

**Supplementary Table S2:** Expression Matrix AUREO1b-OE VS mVenus.xlsx

**Supplementary Table S3:** Expression Matrix σ70.4-OE VS mVenus.xlsx

**Supplementary Table S4:** Expression Matrix HSF1-OE VS mVenus.xlsx

**Supplementary Table S5:** Expression Matrix HSF3-OE VS mVenus.xlsx

**Supplementary Table S6:** Genes associated with metabolic pathways.xlsx

**Supplementary File S1:** Sequence of L2-1_p49-AUREO1b_mVenus_ZeoR.gd

**Supplementary File S2:** Sequence of L2-1_p49-HSF1_mVenus_ZeoR.gd

**Supplementary File S3:** Sequence of L2-1_p49-HSF3_mVenus_ZeoR.gd

**Supplementary File S4:** Sequence of L2-1_p49-Sigma70.4_mVenus_ZeoR.gd

**Supplementary File S5:** Sequence of L2-1_p49-mVenus_ZeoR.gd

**Supplementary File S6:** GO molecular function plot in R.r

## 8. Data Availability

The RNA-seq data generated and analyzed in this study have been deposited in the NCBI Sequence Read Archive (SRA) under the accession number PRJNA1299405.

## 9. Declaration of competing interests

The authors declare no conflict of interest.

## Notes

### Competing Interest Statement

The authors have declared no competing interest.

## References

[1] M. Rizwan, G. Mujtaba, S.A. Memon, K. Lee, N. Rashid, Exploring the potential of microalgae for new biotechnology applications and beyond: A review, Renew. Sustain. Energy Rev. 92 (2018) 394–404. 10.1016/j.rser.2018.04.034.

[2] M.T. Russo, A. Rogato, M. Jaubert, B.J. Karas, A. Falciatore, *Phaeodactylum tricornutum*: An established model species for diatom molecular research and an emerging chassis for algal synthetic biology, J. Phycol. 59 (2023) 1114–1122. 10.1111/jpy.13400.

[3] J. George, T. Kahlke, R.M. Abbriano, U. Kuzhiumparambil, P.J. Ralph, M. Fabris, Metabolic Engineering Strategies in Diatoms Reveal Unique Phenotypes and Genetic Configurations With Implications for Algal Genetics and Synthetic Biology, Front. Bioeng. Biotechnol. 8 (2020) 513. 10.3389/fbioe.2020.00513.

[4] F. Pruckner, L. Morelli, P. Patwari, M. Fabris, Remodeling of the terpenoid metabolism during prolonged phosphate depletion in the marine diatom *Phaeodactylum tricornutum*, J. Phycol. 61 (2025) 512–528. 10.1111/jpy.70014.

[5] Y.J. Zhou, Expanding the terpenoid kingdom, Nat. Chem. Biol. 14 (2018) 1069–1070. 10.1038/s41589-018-0167-4.

[6] E.J. Dufourc, Sterols and membrane dynamics, J. Chem. Biol. 1 (2008) 63–77. 10.1007/s12154-008-0010-6.

[7] I. Domonkos, M. Kis, Z. Gombos, B. Ughy, Carotenoids, versatile components of oxygenic photosynthesis, Prog. Lipid Res. 52 (2013) 539–561. 10.1016/j.plipres.2013.07.001.

[8] H.S. Tsui, C.F. Clarke, Ubiquinone Biosynthetic Complexes in Prokaryotes and Eukaryotes, Cell Chem. Biol. 26 (2019) 465–467. 10.1016/j.chembiol.2019.04.005.

[9] W.P. Cochlan, B.D. Bill, A.B. Cailipan, V.L. Trainer, Domoic acid production by Pseudo-nitzschia australis: Re-evaluating the role of macronutrient limitation on toxigenicity, Harmful Algae 125 (2023) 102431. 10.1016/j.hal.2023.102431.

[10] C. Gallo, G. Nuzzo, G. d’Ippolito, E. Manzo, A. Sardo, A. Fontana, Sterol Sulfates and Sulfotransferases in Marine Diatoms, in: Methods Enzymol., Elsevier, 2018: pp. 101–138. 10.1016/bs.mie.2018.03.003.

[11] E. Carsanba, M. Pintado, C. Oliveira, Fermentation Strategies for Production of Pharmaceutical Terpenoids in Engineered Yeast, Pharmaceuticals 14 (2021) 295. 10.3390/ph14040295.

[12] M. Bertrand, Carotenoid biosynthesis in diatoms, Photosynth. Res. 106 (2010) 89–102. 10.1007/s11120-010-9589-x.

[13] S. Tian, D. Wang, L. Yang, Z. Zhang, Y. Liu, A systematic review of 1-Deoxy-D-xylulose-5-phosphate synthase in terpenoid biosynthesis in plants, Plant Growth Regul. 96 (2022) 221–235. 10.1007/s10725-021-00784-8.

[14] W.-Y. Gao, H. Li, 1-Deoxy-d-xylulose 5-phosphate reductoisomerase, the first committed enzyme in the MEP terpenoid biosynthetic pathway—Its chemical mechanism and inhibition, in: Metalloenzymes, Elsevier, 2024: pp. 375–390. 10.1016/B978-0-12-823974-2.00026-7.

[15] E. Cordoba, M. Salmi, P. Leon, Unravelling the regulatory mechanisms that modulate the MEP pathway in higher plants, J. Exp. Bot. 60 (2009) 2933–2943. 10.1093/jxb/erp190.

[16] A. Athanasakoglou, S.C. Kampranis, Diatom isoprenoids: Advances and biotechnological potential, Biotechnol. Adv. 37 (2019) 107417. 10.1016/j.biotechadv.2019.107417.

[17] W. Ding, Y. Ye, L. Yu, M. Liu, J. Liu, Physiochemical and molecular responses of the diatom Phaeodactylum tricornutum to illumination transitions, Biotechnol. Biofuels Bioprod. 16 (2023) 103. 10.1186/s13068-023-02352-w.

[18] K.J. Lauersen, Eukaryotic microalgae as hosts for light-driven heterologous isoprenoid production, Planta 249 (2019) 155–180. 10.1007/s00425-018-3048-x.

[19] J. Pollier, E. Vancaester, U. Kuzhiumparambil, C.E. Vickers, K. Vandepoele, A. Goossens, M. Fabris, A widespread alternative squalene epoxidase participates in eukaryote steroid biosynthesis, Nat. Microbiol. 4 (2019) 226–233. 10.1038/s41564-018-0305-5.

[20] A.C. Jaramillo-Madrid, R. Abbriano, J. Ashworth, M. Fabris, M. Pernice, P.J. Ralph, Overexpression of Key Sterol Pathway Enzymes in Two Model Marine Diatoms Alters Sterol Profiles in Phaeodactylum tricornutum, Pharmaceuticals 13 (2020) 481. 10.3390/ph13120481.

[21] A.C. Jaramillo-Madrid, J. Ashworth, M. Fabris, P.J. Ralph, The unique sterol biosynthesis pathway of three model diatoms consists of a conserved core and diversified endpoints, Algal Res. 48 (2020) 101902. 10.1016/j.algal.2020.101902.

[22] M.D. Engstrom, B.F. Pfleger, Transcription control engineering and applications in synthetic biology, Synth. Syst. Biotechnol. 2 (2017) 176–191. 10.1016/j.synbio.2017.09.003.

[23] J. Song, H. Zhao, L. Zhang, Z. Li, J. Han, C. Zhou, J. Xu, X. Li, X. Yan, The Heat Shock Transcription Factor PtHSF1 Mediates Triacylglycerol and Fucoxanthin Synthesis by Regulating the Expression of *GPAT3* and *DXS* in *Phaeodactylum tricornutum*, Plant Cell Physiol. 64 (2023) 622–636. 10.1093/pcp/pcad023.

[24] H. Zhao, Y. Liu, Z. Zhu, Q. Feng, Y. Ye, J. Zhang, J. Han, C. Zhou, J. Xu, X. Yan, X. Li, Mediator subunit MED8 interacts with heat shock transcription factor HSF3 to promote fucoxanthin synthesis in the diatom *Phaeodactylum tricornutum*, New Phytol. 241 (2024) 1574–1591. 10.1111/nph.19467.

[25] O. Ait-Mohamed, A.M.G. Novák Vanclová, N. Joli, Y. Liang, X. Zhao, A. Genovesio, L. Tirichine, C. Bowler, R.G. Dorrell, PhaeoNet: A Holistic RNAseq-Based Portrait of Transcriptional Coordination in the Model Diatom Phaeodactylum tricornutum, Front. Plant Sci. 11 (2020) 590949. 10.3389/fpls.2020.590949.

[26] J.A. Berges, D.J. Franklin, P.J. Harrison, EVOLUTION OF AN ARTIFICIAL SEAWATER MEDIUM: IMPROVEMENTS IN ENRICHED SEAWATER, ARTIFICIAL WATER OVER THE LAST TWO DECADES, J. Phycol. 37 (2001) 1138–1145. 10.1046/j.1529-8817.2001.01052.x.

[27] B. Pollak, T. Matute, I. Nuñez, A. Cerda, C. Lopez, V. Vargas, A. Kan, V. Bielinski, P. Von Dassow, C.L. Dupont, F. Federici, Universal loop assembly: open, efficient and cross-kingdom DNA fabrication, Synth. Biol. 5 (2020) ysaa001. 10.1093/synbio/ysaa001.

28. Ensembl Protists, (n.d.). http://protists.ensembl.org/index.html (accessed April 13, 2026).

29. Cloud-based platform for biotech R&D | Benchling, (n.d.). https://www.benchling.com (accessed April 13, 2026).

[30] B.J. Karas, R.E. Diner, S.C. Lefebvre, J. McQuaid, A.P.R. Phillips, C.M. Noddings, J.K. Brunson, R.E. Valas, T.J. Deerinck, J. Jablanovic, J.T.F. Gillard, K. Beeri, M.H. Ellisman, J.I. Glass, C.A. Hutchison Iii, H.O. Smith, J.C. Venter, A.E. Allen, C.L. Dupont, P.D. Weyman, Designer diatom episomes delivered by bacterial conjugation, Nat. Commun. 6 (2015) 6925. 10.1038/ncomms7925.

[31] Protein BLAST: search protein databases using a protein query, (n.d.). https://blast.ncbi.nlm.nih.gov/Blast.cgi?PAGE=Proteins (accessed April 13, 2026).

[32] NGPhylogeny.fr, (n.d.). https://ngphylogeny.fr/ (accessed April 13, 2026).

[33] I. Letunic, P. Bork, Interactive Tree of Life (iTOL) v6: recent updates to the phylogenetic tree display and annotation tool, Nucleic Acids Res. 52 (2024) W78–W82. 10.1093/nar/gkae268.

[34] S.N. Coesel, More than a photoreceptor: aureochromes are intrinsic to the diatom light-regulated transcriptional network, J. Exp. Bot. 75 (2024) 1786–1790. 10.1093/jxb/erae004.

[35] M.J.J. Huysman, A.E. Fortunato, M. Matthijs, B.S. Costa, R. Vanderhaeghen, H. Van Den Daele, M. Sachse, D. Inzé, C. Bowler, P.G. Kroth, C. Wilhelm, A. Falciatore, W. Vyverman, L. De Veylder, AUREOCHROME1a-Mediated Induction of the Diatom-Specific Cyclin *dsCYC2* Controls the Onset of Cell Division in Diatoms (*Phaeodactylum tricornutum*), Plant Cell 25 (2013) 215–228. 10.1105/tpc.112.106377.

[36] B. Schellenberger Costa, M. Sachse, A. Jungandreas, C.R. Bartulos, A. Gruber, T. Jakob, P.G. Kroth, C. Wilhelm, Aureochrome 1a Is Involved in the Photoacclimation of the Diatom Phaeodactylum tricornutum, PLoS ONE 8 (2013) e74451. 10.1371/journal.pone.0074451.

[37] S.H. Im, B. Lepetit, N. Mosesso, S. Shrestha, L. Weiss, M. Nymark, R. Roellig, C. Wilhelm, E. Isono, P.G. Kroth, Identification of promoter targets by Aureochrome 1a in the diatom *Phaeodactylum tricornutum*, J. Exp. Bot. 75 (2024) 1834–1851. 10.1093/jxb/erad478.

[38] E. Herman, M. Sachse, P.G. Kroth, T. Kottke, Blue-Light-Induced Unfolding of the Jα Helix Allows for the Dimerization of Aureochrome-LOV from the Diatom *Phaeodactylum tricornutum*, Biochemistry 52 (2013) 3094–3101. 10.1021/bi400197u.

[39] E. Herman, T. Kottke, Allosterically Regulated Unfolding of the A′α Helix Exposes the Dimerization Site of the Blue-Light-Sensing Aureochrome-LOV Domain, Biochemistry 54 (2015) 1484–1492. 10.1021/bi501509z.

[40] H. Zhang, X. Xiong, K. Guo, M. Zheng, T. Cao, Y. Yang, J. Song, J. Cen, J. Zhang, Y. Jiang, S. Feng, L. Tian, X. Li, A rapid aureochrome opto-switch enables diatom acclimation to dynamic light, Nat. Commun. 15 (2024) 5578. 10.1038/s41467-024-49991-7.

[41] M. Mann, M. Serif, T. Jakob, P.G. Kroth, C. Wilhelm, PtAUREO1a and PtAUREO1b knockout mutants of the diatom Phaeodactylum tricornutum are blocked in photoacclimation to blue light, J. Plant Physiol. 217 (2017) 44–48. 10.1016/j.jplph.2017.05.020.

[42] M.S. Paget, J.D. Helmann, The σ70family of sigma factors, Genome Biol. 4 (2003) 203. 10.1186/gb-2003-4-1-203.

[43] L.A. Allison, The role of sigma factors in plastid transcription*, Biochimie 82 (2000) 537–548. 10.1016/S0300-9084(00)00611-8.

[44] Y. Lin, J. Feng, H. Fang, W. Huang, K. Guo, X. Liu, S. Wang, X. Liu, The Expression Characteristics and Potential Functions of Heat Shock Factors in Diatom *Phaeodactylum tricornutum*, Phyton-Int. J. Exp. Bot. 93 (2024) 2583–2596. 10.32604/phyton.2024.055616.

[45] Y. Su, J. Hu, M. Xia, J. Chen, W. Meng, C. Qian, Y. Shu, C. Wang, X. Wang, K. Salehi-Ashtiani, S. Brynjólfsson, J. Lin, Y. Li, H. Zhang, L. Wang, W. Fu, An undiscovered circadian clock to regulate phytoplankton photosynthesis, PNAS Nexus 3 (2024) pgae497. 10.1093/pnasnexus/pgae497.

[46] A. Banerjee, E. Herman, M. Serif, M. Maestre-Reyna, S. Hepp, R. Pokorny, P.G. Kroth, L.-O. Essen, T. Kottke, Allosteric communication between DNA-binding and light-responsive domains of diatom class I aureochromes, Nucleic Acids Res. 44 (2016) 5957–5970. 10.1093/nar/gkw420.

[47] J.M. Buck, J. Sherman, C.R. Bártulos, M. Serif, M. Halder, J. Henkel, A. Falciatore, J. Lavaud, M.Y. Gorbunov, P.G. Kroth, P.G. Falkowski, B. Lepetit, Lhcx proteins provide photoprotection via thermal dissipation of absorbed light in the diatom Phaeodactylum tricornutum, Nat. Commun. 10 (2019) 4167. 10.1038/s41467-019-12043-6.

[48] T.-B. Hao, T. Jiang, H.-P. Dong, L. Ou, X. He, Y.-F. Yang, Light-harvesting protein Lhcx3 is essential for high light acclimation of Phaeodactylum tricornutum, AMB Express 8 (2018) 174. 10.1186/s13568-018-0703-3.

[49] N. Shalygo, O. Czarnecki, E. Peter, B. Grimm, Expression of chlorophyll synthase is also involved in feedback-control of chlorophyll biosynthesis, Plant Mol. Biol. 71 (2009) 425–436. 10.1007/s11103-009-9532-8.

[50] M. Plöchinger, S. Schwenkert, L. Von Sydow, W.P. Schröder, J. Meurer, Functional Update of the Auxiliary Proteins PsbW, PsbY, HCF136, PsbN, TerC and ALB3 in Maintenance and Assembly of PSII, Front. Plant Sci. 7 (2016). 10.3389/fpls.2016.00423.

[51] S. Uto, K. Kawakami, Y. Umena, M. Iwai, M. Ikeuchi, J.-R. Shen, N. Kamiya, Mutual relationships between structural and functional changes in a PsbM-deletion mutant of photosystem II, Faraday Discuss. 198 (2017) 107–120. 10.1039/C6FD00213G.

[52] E. Rayko, F. Maumus, U. Maheswari, K. Jabbari, C. Bowler, Transcription factor families inferred from genome sequences of photosynthetic stramenopiles, New Phytol. 188 (2010) 52–66. 10.1111/j.1469-8137.2010.03371.x.

[53] M. Chenge-Espinosa, E. Cordoba, C. Romero-Guido, G. Toledo-Ortiz, P. León, Shedding light on the methylerythritol phosphate (MEP)-pathway: long hypocotyl 5 (HY5)/phytochrome-interacting factors (PIFs) transcription factors modulating key limiting steps, Plant J. 96 (2018) 828–841. 10.1111/tpj.14071.

[54] K. Ameztoy, Á.M. Sánchez-López, F.J. Muñoz, A. Bahaji, G. Almagro, E. Baroja-Fernández, S. Gámez-Arcas, N. De Diego, K. Doležal, O. Novák, A. Pěnčík, A. Alpízar, M. Rodríguez-Concepción, J. Pozueta-Romero, Proteostatic Regulation of MEP and Shikimate Pathways by Redox-Activated Photosynthesis Signaling in Plants Exposed to Small Fungal Volatiles, Front. Plant Sci. 12 (2021). 10.3389/fpls.2021.637976.

[55] A. Banerjee, Y. Wu, R. Banerjee, Y. Li, H. Yan, T.D. Sharkey, Feedback Inhibition of Deoxy-d-xylulose-5-phosphate Synthase Regulates the Methylerythritol 4-Phosphate Pathway *, J. Biol. Chem. 288 (2013) 16926–16936. 10.1074/jbc.M113.464636.

[56] C.A. Schuhr, T. Radykewicz, S. Sagner, C. Latzel, M.H. Zenk, D. Arigoni, A. Bacher, F. Rohdich, W. Eisenreich, Quantitative assessment of crosstalk between the two isoprenoid biosynthesis pathways in plants by NMR spectroscopy, Phytochem. Rev. 2 (2003) 3–16. 10.1023/B:PHYT.0000004180.25066.62.

[57] O. Laule, A. Fürholz, H.-S. Chang, T. Zhu, X. Wang, P.B. Heifetz, W. Gruissem, M. Lange, Crosstalk between cytosolic and plastidial pathways of isoprenoid biosynthesis in Arabidopsis thaliana, Proc. Natl. Acad. Sci. 100 (2003) 6866–6871. 10.1073/pnas.1031755100.

[58] X. Huang, F. Wang, O.U. Rehman, X. Hu, F. Zhu, R. Wang, L. Xu, Y. Cui, S. Huo, Influence of Light Regimes on Production of Beneficial Pigments and Nutrients by Microalgae for Functional Plant-Based Foods, Foods 14 (2025) 2500. 10.3390/foods14142500.

