## Supplementary figures file (S1-S4) for "AUREOCHROME 1b is a transcriptional regulator of pigment biosynthesis in *Phaeodactylum tricornutum*"

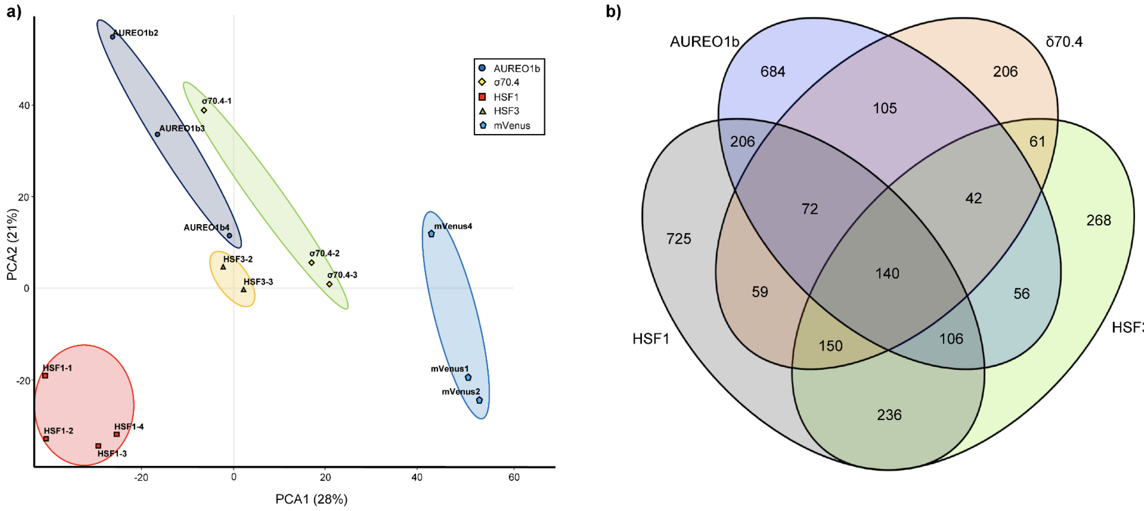


**Figure S1. Quality control and differential expression overview of RNA-seq data.** (a) Principal component analysis of all biological replicates demonstrates separation of individual groups. (b) Venn diagram demonstrating common differentially expressed genes between different transcription factor overexpressing lines.

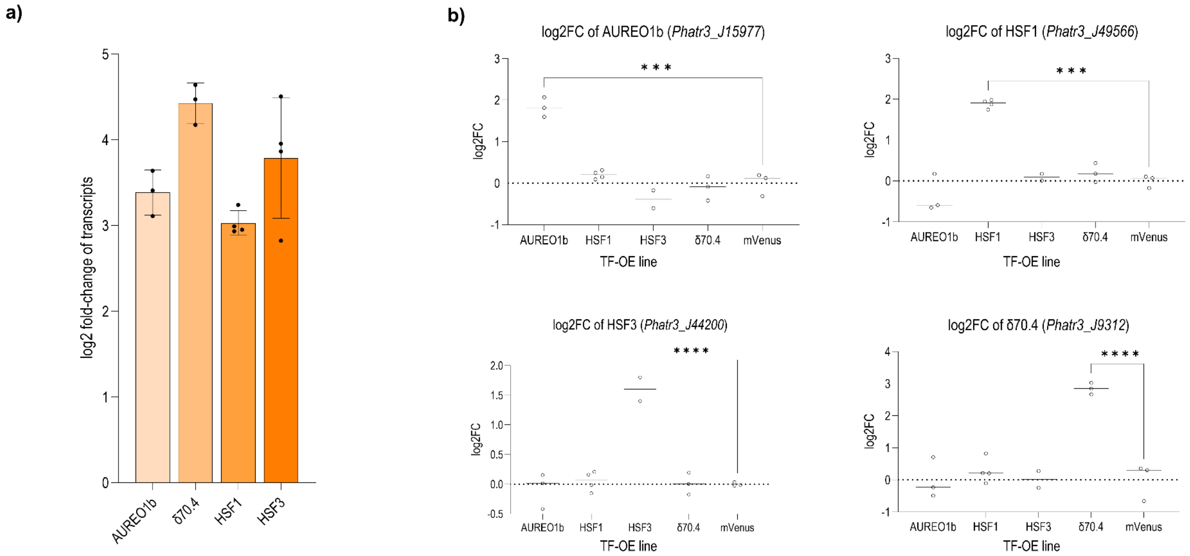


**Figure S2. Validation of transcription factor (TF) overexpression.** (a) Quantitative PCR (qPCR) confirmation of overexpression: for each TF-OE line, the bar height shows the fold-increase in that transcription factor's mRNA relative to the mVenus control. (b) RNA-seq derived expression change of the transcription factor in different TF-overexpressing (TF-OE) lines (x-axis), as relative change to mVenus lines, are plotted in different graphs for AUREO1b, HSF1, HSF3, and σ70.4. Significant differences between groups were determined by Tukey-test ($p$-value < 0.01) and are indicated by asterisks.

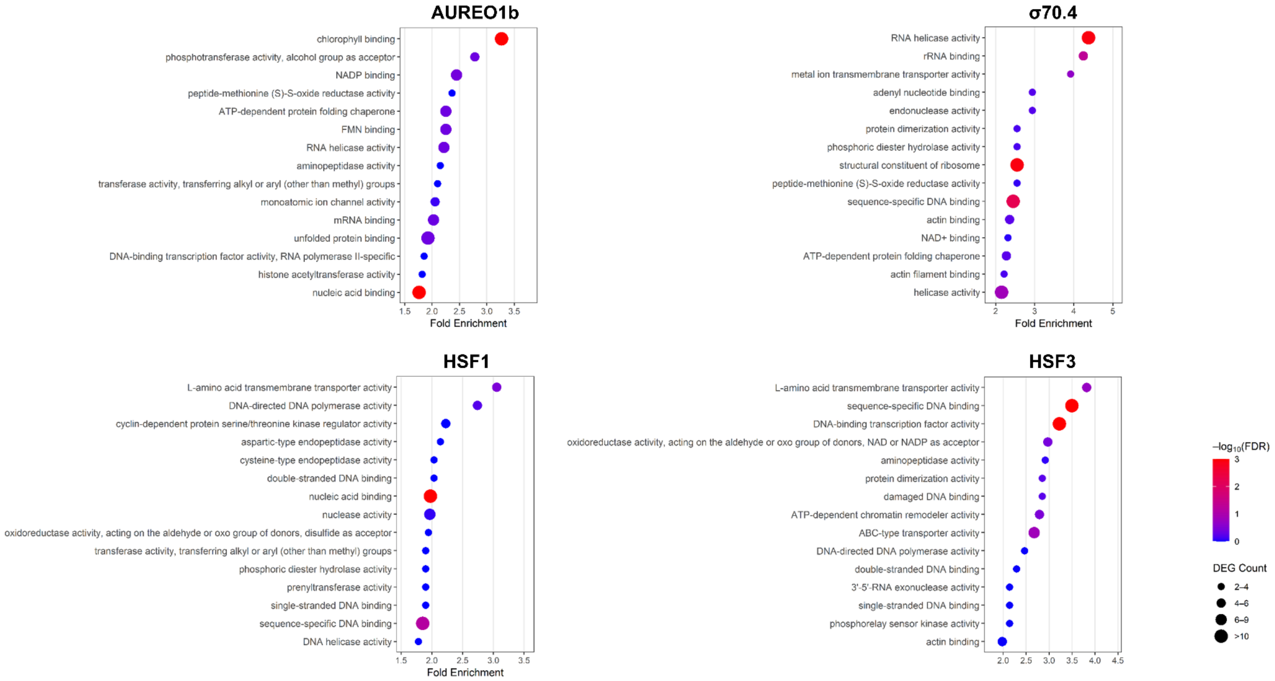


**Figure S3. Gene ontology (GO) analysis of enriched molecular functions of differentially expressed genes of transcription factor (TF) overexpressing (TF-OE) lines.** The colour of the circle indicates the false discovery rate, with red indicating high confidence, and blue indicating low confidence. Circle size indicates the number of genes of a certain molecular function that were among the differentially expressed genes of a transcription factor. Bigger circles indicate more genes.

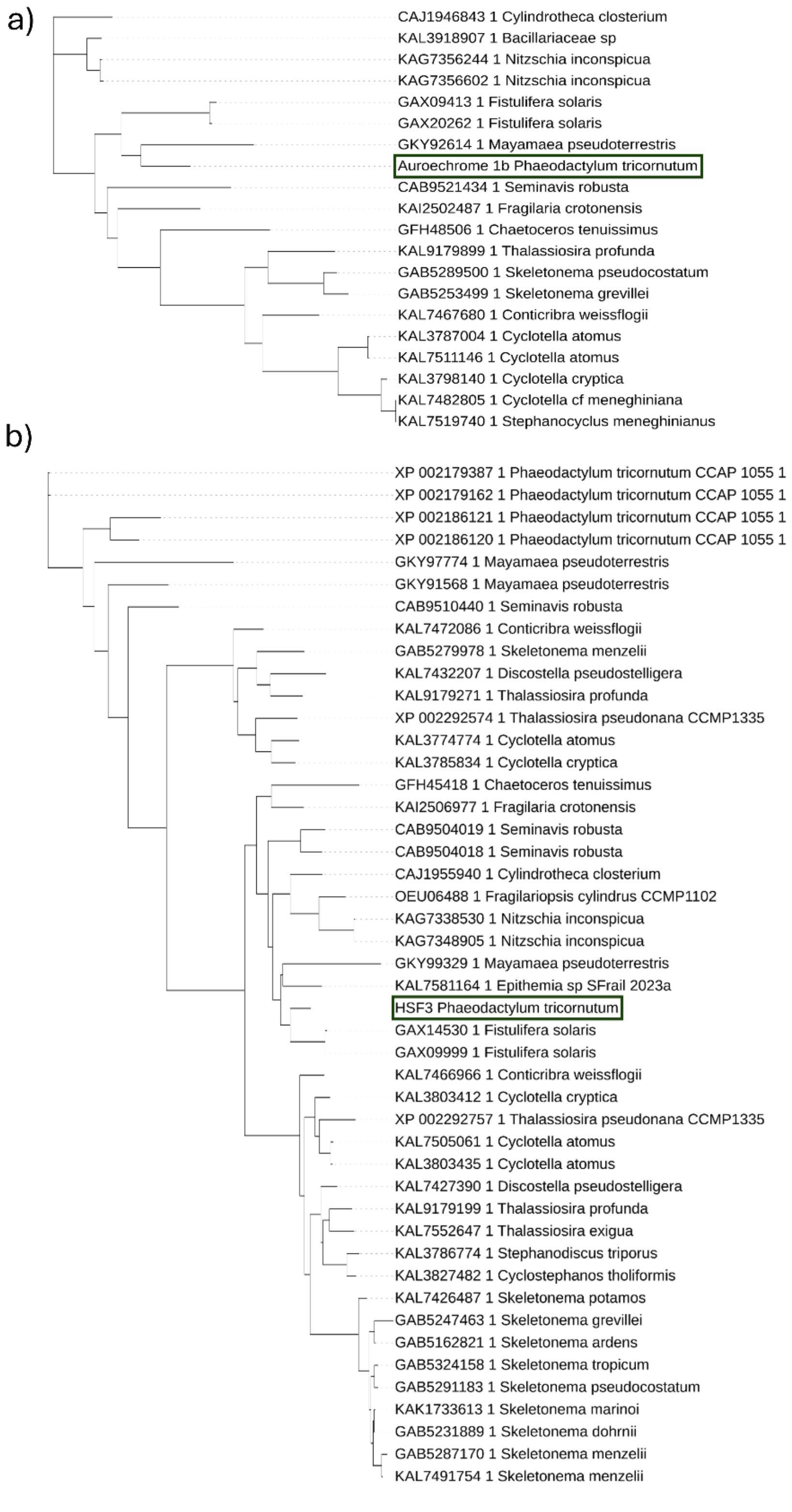


**Figure S4. Phylogenetic analysis of proteins homologous to (a) Aureochrome 1b and (b) Heat Shock Factor 3 (HSF3).** Homologs were identified with NCBI BLASTP (E-value < 0.01; query coverage ≥ 75%). BLAST hits were analyzed using NGPhylogeny.fr with default parameters, and tree graphics were formatted in iTOL (see methods). Original blast query input is indicated as black box.
