## Supplementary files S1-S6 for "AUREOCHROME 1b is a transcriptional regulator of pigment biosynthesis in *Phaeodactylum tricornutum*": File S6_GO molecular function plot.R

```
# If you haven’t installed these yet, do so here:
if (!requireNamespace("BiocManager", quietly=TRUE)) {
  install.packages("BiocManager")
}
# On CRAN:
install.packages(c("ggplot2", "dplyr", "tidyr"))

BiocManager::install(c("clusterProfiler", "biomaRt"))


# Then load them:
library(dplyr)
library(tidyr)
library(ggplot2)
library(clusterProfiler)
library(biomaRt)
library(grid)      # for unit()
library(gtable)    # for gtable manipulation
library(scales)    # for squish()
#### Set WD and load your gene names
# Example gene list


directory_fetchfiles <- "//tek/dfs/data/TEK-FD1/AlgaeSynBio/Lab Members/Florian/Code/output"
directory_dropfiles <- "//tek/dfs/data/TEK-FD1/AlgaeSynBio/Lab Members/Florian/Code/output"
setwd(directory_fetchfiles)

#to loop through all text files in a folder
txtfiles <-  list.files(pattern     = "\\.txt$",
                        full.names  = FALSE,
                        ignore.case = TRUE)

for (txtfile in txtfiles) {

  
# Rest of the code
inputfilename <- txtfile
setwd(directory_fetchfiles)  


genes_df <- read.delim(inputfilename, stringsAsFactors = FALSE)
colnames(genes_df) <- "GeneID"

# Extract the vector of character IDs:
genes <- genes_df$GeneID


#### 2) Query Ensembl Protists for all <gene → GOID> pairs, then filter to MF only

# (2a) Connect to Ensembl Protists
mart <- useMart(
  biomart = "protists_mart",
  host    = "https://protists.ensembl.org"
)

# (2b) Select the P. tricornutum dataset
mart_pt <- useDataset("ptricornutum_eg_gene", mart = mart)

# (2c) Get all (ensembl_gene_id, go_id, namespace_1003, name_1006)
#      where namespace_1003 == "molecular_function"
go_data_raw <- getBM(
  attributes = c("ensembl_gene_id","go_id","namespace_1003","name_1006"),
  mart       = mart_pt
) %>%
  dplyr::filter(
    !is.na(go_id),
    go_id != "",
    namespace_1003 == "molecular_function"
  ) %>%
  dplyr::select(ensembl_gene_id, go_id, name_1006) %>%
  dplyr::distinct()

# (2d) Build TERM2GENE_MF:  
#      a two‐column DF where col1 = GO ID, col2 = EnsemblGeneID
TERM2GENE_MF <- go_data_raw %>%
  dplyr::select(term = go_id, gene = ensembl_gene_id) %>%
  distinct()

TERM2NAME_MF <- go_data_raw %>%
  dplyr::select(term = go_id, term_name = name_1006) %>%
  distinct()

# Quick check:
head(TERM2GENE_MF)
#        term           gene
# 1  GO:0003677  Phatr3_0000123
# 2  GO:0003700  Phatr3_0000456
# …  

head(TERM2NAME_MF)
#        term                      term_name
# 1  GO:0003677      DNA binding
# 2  GO:0003700 transcription factor activity, sequence-specific DNA binding
# …


#### 4) Run GO‐MF enrichment (ORA) with clusterProfiler::enricher()

ego_mf <- enricher(
  gene       = genes,
  TERM2GENE  = TERM2GENE_MF,
  TERM2NAME  = TERM2NAME_MF,     # <- new addition
  universe = unique(TERM2GENE_MF$gene),
  pAdjustMethod = "BH",
  qvalueCutoff  = 0.05
)

# Check the result:
head(ego_mf@result)


#### 5) Compute Fold Enrichment & –log₁₀(FDR), then pick the top 15 terms

# 5a) Extract `ego_mf@result` into a data.frame
res <- as.data.frame(ego_mf@result)

# 5b) Parse “GeneRatio” and “BgRatio” strings into numerators/denominators
res2 <- res %>%
  separate(GeneRatio, into = c("Count_in_list", "Size_list"), sep = "/", convert = TRUE)  %>%
  separate(BgRatio,   into = c("Count_in_bg",   "Size_bg"),   sep = "/", convert = TRUE)  %>%
  mutate(
    # 
    # FoldEnrichment = (Count_in_list / Size_list) / (Count_in_bg / Size_bg)
    FoldEnrichment = (Count_in_list / Size_list) / (Count_in_bg / Size_bg),
    logFDR         = -log10(p.adjust)  # so that a small FDR → large logFDR → “hot” color
  )

# 5c) Sort by descending FoldEnrichment and keep the top 15 rows
top15 <- res2 %>%
  arrange(desc(FoldEnrichment)) %>%
  slice(1:7)

# SECTION 6 (FINAL): Fixed legends and fixed 4″×4″ panel using gtable
#                   • Color gradient (–log10(FDR)) from 0 → 3
#                   • DEG count binned into: 1, 2–4, 4–6, 6–9, >10
#                   • Highest FoldEnrichment term at the top
#                   • No y‐axis label
# ──────────────────────────────────────────────────────────────────────────────


# 1) Reorder 'Description' so the highest FoldEnrichment term is drawn at the top:
top15 <- top15 %>%
  mutate(
    Description = factor(
      Description,
      levels = unique(Description[order(FoldEnrichment)])
      # order(FoldEnrichment) gives indices from lowest→highest,
      # so the last level corresponds to highest FE and is drawn at the top.
    )
  )

# 2) Bin 'Count_in_list' into fixed categories for the size legend:
top15 <- top15 %>%
  mutate(
    sizeBin = cut(
      Count_in_list,
      breaks = c(-Inf, 1, 4, 6, 9, Inf),
      labels = c("1", "2–4", "4–6", "6–9", ">10"),
      right  = TRUE
    )
  )

# 3) Determine how many MF categories we actually have:
n_terms <- nrow(top15)  # could be fewer than 15

# 4) Compute the panel height in inches so that spacing per term = (4" / 15):
base_panel_height <- 4  # inches when 15 terms
panel_height_in   <- base_panel_height * n_terms / 15

# Panel width remains fixed at 4 inches:
panel_width_in <- 2

# 5) Build the ggplot (no aspect.ratio; we’ll override panel size via gtable):
x_min <- min(top15$FoldEnrichment)*0.8
x_max <- max(top15$FoldEnrichment)*1.2

p <- ggplot(top15, aes(x = FoldEnrichment, y = Description)) +
  geom_point(
    aes(
      size  = sizeBin,       # discrete size categories
      color = logFDR         # continuous –log10(FDR)
    )
  ) +
  # 5a) Fix x‐axis range (same for all plots)
  scale_x_continuous(
    limits = c(x_min, x_max),
    expand = c(0, 0)
  ) +
  # 5b) Color gradient: 0→3, values outside squished to nearest limit
  scale_color_gradient(
    low    = "blue",
    high   = "red",
    limits = c(0, 3),
    oob    = squish,
    breaks = c(0, 1, 2, 3),
    labels = c("0", "1", "2", "3")
  ) +
  # 5c) Fixed size mapping for the five bins
  scale_size_manual(
    values = c(
      "1"   = 2,
      "2–4" = 3,
      "4–6" = 4,
      "6–9" = 5,
      ">10" = 6
    ),
    breaks = c("1", "2–4", "4–6", "6–9", ">10"),
    labels = c("1", "2–4", "4–6", "6–9", ">10")
  ) +
  # 5d) Remove y‐axis label, set other labels
  labs(
    x     = "Fold Enrichment",
    y     = NULL,
    color = expression(paste("–log"[10], "(FDR)")),
    size  = "DEG Count"
  ) +
  ggtitle("GO:MF enrichment for P. tricornutum DEGs\n(top terms by FoldEnrichment)") +
  theme_bw() +
  theme(
    panel.grid.major.y = element_blank(),
    panel.grid.minor   = element_blank(),
    plot.title         = element_text(hjust = 0.5),
    axis.text.y        = element_text(size = 10)
  )

# 6) Convert ggplot to a gtable
g <- ggplotGrob(p)

# 7) Find the panel’s location in the layout
panel_index <- which(g$layout$name == "panel")

# 8) Override the panel’s width and height to the computed dimensions:
g$widths[g$layout[panel_index, ]$l]  <- unit(panel_width_in, "in")
g$heights[g$layout[panel_index, ]$t] <- unit(panel_height_in, "in")

# 9) Save to a fixed‐width PNG. We’ll keep overall device height at 6″,
#    so panel appears shorter when fewer than 15 terms. Leftover space becomes margins.
output_name <- paste0(tools::file_path_sans_ext(basename(inputfilename)), ".png")
png(
  filename = output_name,
  width    = 12,      # total device width in inches
  height   = 12,      # total device height in inches
  units    = "in",
  res      = 600
)
grid.draw(g)
dev.off()

}
```
